# Scalability of Random Forest in Myoelectric Control

**DOI:** 10.1101/2025.03.04.641468

**Authors:** Xinyu Jiang, Chenfei Ma, Kianoush Nazarpour

**Affiliations:** School of Informatics, The University of Edinburgh, Edinburgh, United Kingdom

**Keywords:** EMG, myoelectric control, scalable machine learning, random forest

## Abstract

**Objective:** Myoelectric control systems translate electromyographic (EMG) signals into control commands, enabling immersive human-robot interactions in the real world and the Metaverse. The variability of EMG due to various confounding factors leads to significant performance degradation. Such variability can be mitigated by training a highly generalisable but massively parameterized deep neural network, which can be effectively scaled using a vast dataset. We aim to find an alternative simple, explainable, efficient and parallelisable model, which can flexibly scale up with a larger dataset and scale down to reduce model size, will significantly facilitate the practical implementation of myoelectric control.

**Approach:** In this work, we discuss the scalability of a random forest (RF) for myoelectric control. We show how to scale an RF up and down during the process of pre-training, fine-tuning, and automatic self-calibration. The effects of diverse factors such as bootstrapping, decision tree editing (pre-training, pruning, grafting, appending), and the size of training data are systematically studied using EMG data from 106 participants including both low- and high-density electrodes.

**Main results:** We examined several factors that affect the size and accuracy of the model. The best solution could reduce the size of RF models by *≈*500 *×*, with the accuracy reduced by only 1.5%. Importantly, for the first time we report the unique merit of RF that with more EMG electrodes (higher input dimension), the RF model size would be reduced, contrasting all other models.

**Significance:** All of these findings unlock the full potential of RF in real-world applications.

## 1. Introduction

Myoelectric control systems translate electromyographic signals (EMG) under different hand gestures [1, 2, 3] into the control commands of various robotics such as exoskeletons and bionic hands, allowing intuitive and immersive human-robot interactions in the real world and the Metaverse [4, 5, 6, 7, 8, 9, 10, 11, 12]. However, the large variations of EMG due to various confounding factors such as arm positions [13, 14], electrode shift [15], variation of user behavior [16], variation between users [17, 18] and variation between days [19] of EMG characteristics lead to significant performance degradation in real-world applications. Such variabilities can be mitigated by training a highly generalisable but massively parameterized deep neural network, which can be effectively scaled using a vast dataset. In particular, the scaling law of large neural networks has contributed to highly generalisable but extremely large backbone models in both natural language processing [20] and computer vision fields [21]. A very recent study by Meta has also introduced a comprehensive generic deep network-based myoelectric control model [22], which was trained using data from 6527 participants and consisted of a total of 60.2 million parameters. Such large generic neural networks show impressive performance on various myoelectric control tasks.

However, training large neural networks using backpropagation requires extensive datasets and significant computing infrastructure. While these pre-trained networks could achieve impressive results, their intricate design often makes them function as a black box and requires extensive computation resources. Considering that myoelectric control models are generally applied in mobile scenarios, it is crucial to deploy models in compact and wearable systems with limited computational and battery resources. A large neural network-based myoelectric control model with 60.2 million parameters [22] can hardly be embedded in low-cost mobile computing devices, for example microcontrollers. Moreover, for human-centered applications, especially biomedical applications, the used model is expected to be explainable [23].

Most conventional machine learning models other than neural networks are generally considered to be trainable only from scratch, struggling to efficiently handle the significant and progressive increase in training data. A series of our recent studies [24, 25] proved that standard random forest (RF) models are pre-trainable and can be personalized via one-shot fine-tuning. Furthermore, a self-calibrating RF can automatically update the model parameters in an unsupervised way to progressively improve the performance of the model [25] and is highly robust against confounders such as arm positions [13]. Our previous study has also shown that RF-based models are explicable and resistant to electrode corruption [23]. These studies have shown the promising prospects of RF as an alternative in myoelectric control.

In this work, our aim is to further unlock the full potentials of RF in myoelectric control. By revisiting our previously proposed RF-based myoelectric control models, we dive into the scalability of RF and show how to scale up and down a RF during the pre-training, fine-tuning, and automatic self-calibration of RF. We analyzed the effects of various factors such as bootstrapping, decision tree pre-training, pruning, grafting, appending, and the size of training data, through extensive experiments on EMG from 106 participants including both low-density (LD) and high-density (HD) electrodes. We explored several factors affecting the model size and accuracy, to find a solution to trade off between model size and accuracy. Our analyses shows, the size of the RF models can be compressed by ≈ 500× with negligible reduction in accuracy (1.5%). Importantly, we found a unique and promising property of RF that, with more EMG electrodes (higher input dimension) the RF model size would even be reduced. The conclusion of our analyses can hopefully generalise across other tree-based myoelectric control models (e.g., XGBoost), allowing scalability.

## 2. Data Collection

### 2.1. LD Datasets

All participants provided their signed informed consent forms prior to participating in the study. Our experiment was approved by the local ethics committee of the University of Edinburgh (ref: 2019/89177). Data from 86 participants were collected in 4 experiments with the same electrode configuration and involved hand gestures presented in Fig. 1. Specifically, data recorded in 8 Delsys Trigno sensors with a 10–500 Hz passband and a 2000 Hz sampling rate were used in our analyses. A summary of all datasets used in our analyses is presented in Table 1. Detailed descriptions of these data sets can be found in [24, 25, 13]. Here we only provide a brief description.

**Table 1.** Summary of all datasets used in our analyses.

| Dataset | Number of participants | Number of channels | Number of hand gestures | Number of trials | Valid duration per trial (s) | Conditions |
| --- | --- | --- | --- | --- | --- | --- |
| LD dataset 1 | 20 | 8 | 6 | 10 | 4 | Same day & same arm position |
| LD dataset 2 | 18 | 8 | 6 | 51 | 1 | 2 days & same arm position |
| LD dataset 3 | 28 | 8 | 6 | 51 | 1 | 2 days & same arm position |
| LD dataset 4 | 20 | 8 | 6 | 56 | 1 | Same-day & 5 arm positions |
| HD dataset | 20 | 256 | 10 | 12 | 0.5 | 2 days & same arm position |

**Figure 1.**
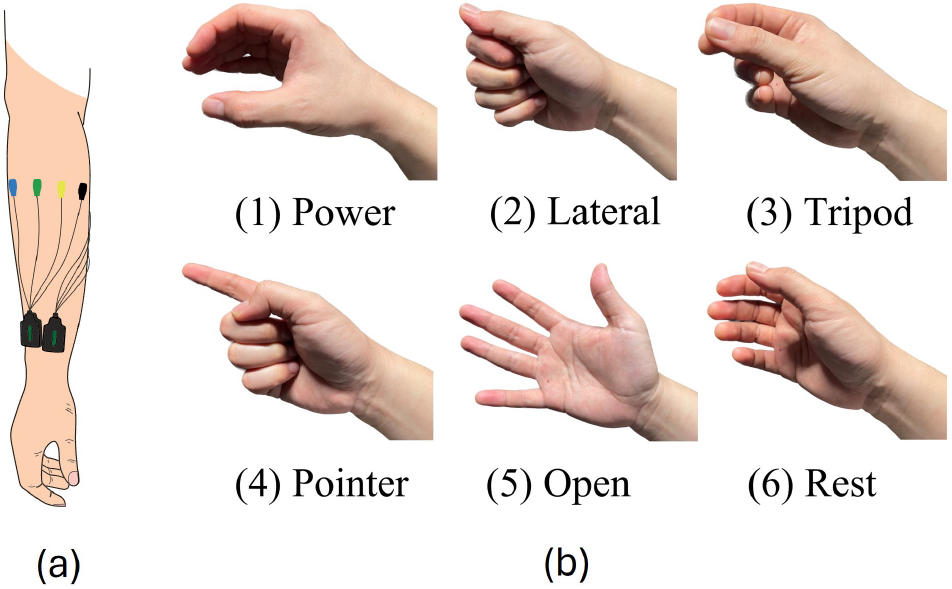
Electrode positions and hand gestures involved in our experiment. This figure was adopted from our previous open access paper [24] under the Creative Commons License (https://creativecommons.org/licenses/by/4.0/).

#### 2.1.1. LD Dataset 1

LD dataset 1 consisted of data from 20 participants (aged 22–43, 12 males, 8 females). For each participant, data from 10 trials were collected for each hand gesture (4 s valid durations per trial) on the same day.

#### 2.1.2. LD Dataset 2

LD dataset 2 consisted of data from 18 *new* participants (aged 22-28, 11 males, 7 females). Each participant performed one trial (with 1 s valid duration used in our analyses) per hand gesture in an initial calibration session. Data collected in the calibration session were used to perform one-shot model training/calibration in our following analyses. The participants then went through a testing session comprising five testing blocks with five tests per hand gesture in each block. The next consecutive day, participants repeated the same testing session directly without a re-calibration session. Consequently, the inter-day variabilities of EMG were taken into account in this dataset.

#### 2.1.3. LD Dataset 3

LD dataset 3 consisted of 28 *new* participants (aged 21–42, 13 males, 15 females). The data collection experiment of this dataset is similar to that of LD dataset 2. The only difference is that, in each testing block, the number of trials for different hand gestures is not necessarily the same to simulate an unbalanced testing block. However, we still kept the total number of trials in both days balanced for a reliable evaluation of the model performance. All of the above data were collected with the participants’ arms pointing vertically toward the ground.

#### 2.1.4. LD Dataset 4

LD dataset 4 consisted of 20 *new* participants (aged 19–31, 10 males, 10 females). The experiment consisted of a calibration session (one trial per hand gesture) followed and a testing session (with 11 testing blocks). The calibration session was performed with the participants’ arm at position 5 (P5) presented in Fig. 2. Data in different testing blocks were collected at varying arm positions listed in Fig. 2 (details in [13]). Accordingly, the inter-posture variabilities of EMG were taken into account in this dataset.

**Figure 2.**
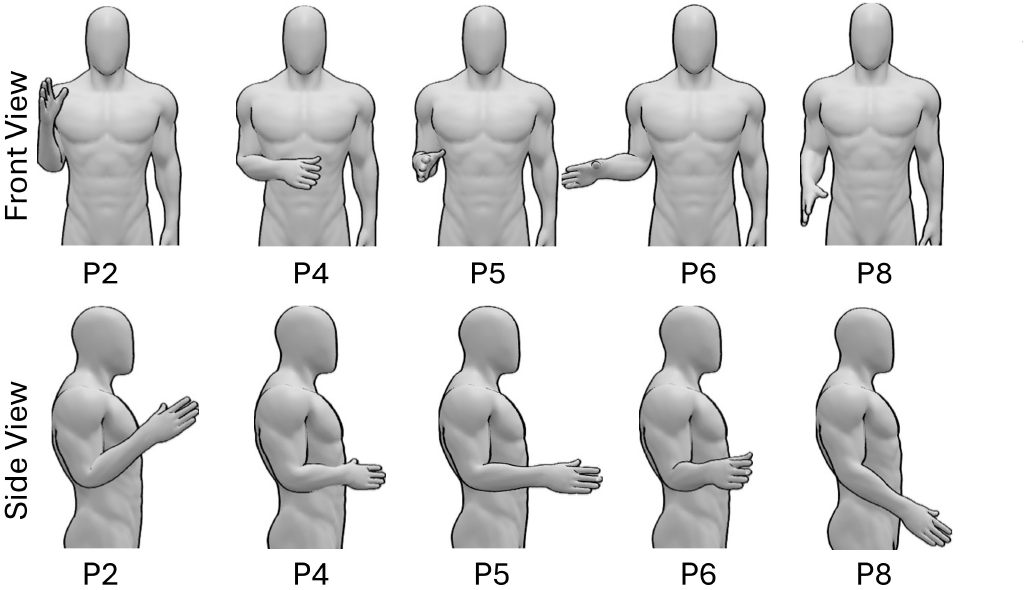
Different arm positions. This figure was adopted from our previous open access paper [13] under the Creative Commons License (https://creativecommons.org/licenses/by/4.0/).

### 2.2. HD Dataset

The open access hyser dataset [26] collected from 20 participants with 256 densely distributed electrodes was used to explore the property of the RF model in HD EMG data. Ten hand gestures commonly used in human-machine interactions were used in our analyzes, as presented in Fig. 3. Participants performed a dynamic hand gesture task within the duration of a trial of 1s. Six trials were repeated for each hand gesture on each day. Data were collected on two days with the same experiment design. The first 500 ms of a trial after the visual cue was viewed as the reaction period of participants, and not involved in the training and testing in our analyzes.

**Figure 3.**
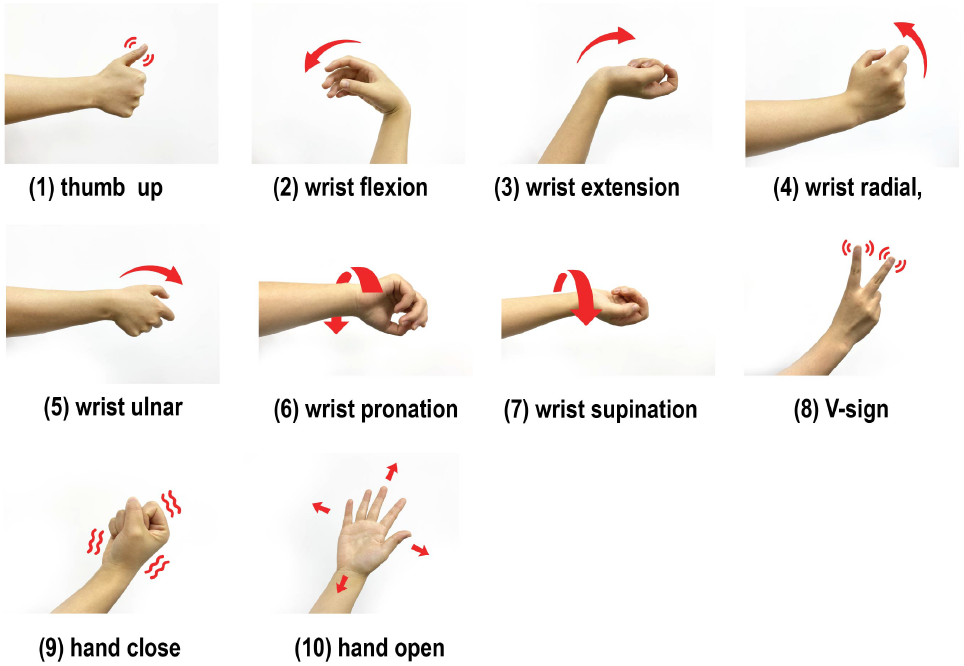
Hand gestures involved in the HD-EMG dataset.

## 3. Methods

### 3.1. Feature Extraction

Ten types of features, namely mean absolute value (MAV), waveform length (WL), root mean square (RMS), slope sign changes (SSC), zero crossings (ZC), peak frequency (PKF), median frequency (MDF), mean frequency (MNF), variance of central frequency (VCF) and skewness (SKW), were extracted from each EMG channel via a sliding window (window length: 200 ms; sliding step: 100 ms). The effectiveness of these features has been solidly validated in previous studies [24, 13].

### 3.2. RF Model Pretraining

The RF model for each target participant was first pre-trained on data from other participants. The pre-trained RF model comprised *N*_*tree*_ decision trees. To train each decision tree, *r* (that is, the ratio of selected samples to the total number of samples) samples were randomly selected (with replacement) by bootstrapping to train each decision tree. The use of only a small proportion of samples in pre-training each decision tree could largely reduce the complexity of each decision tree and encourage diversity among different decision trees.

### 3.3. RF Model Fine-tuning

#### 3.3.1. Decision tree grafting

Decision tree grafting refers to the operation of further growing new branches from the original leaf nodes of a decision tree, so that samples with mixed labels in the same leaf node can be further purified. The new grafted branches were trained using the labeled calibration data.

#### 3.3.2. Decision tree pruning

Decision tree pruning refers to the operation to remove unnecessary branches from a decision tree, so that the original decision logic can be simplified and generalized to a new application setting. The pruning operation was performed using the labeled calibration data as a validation data set to verify if removing a pre-trained branch could contribute to improved model performance.

#### 3.3.3. Decision tree appending

Decision tree appending refers to the operation to implant new decision trees (trained from scratch using labeled calibration data) into the pretrained RF.

More details on above three operations to fine-tune a RF model can be found in our previous work [24]. In this work, we applied and compared the performance of (1) grafting + appending, and (2) pruning + appending strategies.

### 3.4. RF Model Self-calibration

A self-calibrating RF employs a data buffer to save incoming testing samples. Pseudolabels were assigned to these unlabeled test samples jointly by the knowledge learned by the current RF model and the statistical distribution knowledge learned by t-Distributed Stochastic Neighbor Embedding (t-SNE) [27] and K-means clustering. Specifically, all samples in the data buffer were first assigned initial pseudolabels by the predictions of the current RF model (RF-based pseudolabels). Then, all buffered samples were mapped to a 3-dimensional subspace by manifold learning using t-SNE, which is a nonlinear dimensionality reduction technique for embedding high-dimensional data in a low-dimensional space. The distribution of samples in the t-SNE space would be more linear than the original high-dimensional space. Subsequently, K-means clustering was then applied in the t-SNE space to assign the final pseudolabels with the RF-based pseudolabels as the initialization of K-means. The labeled data used in the one-shot calibration and the pseudolabeled data saved in the data buffer were concatenated to train new decision trees and update 40% of the original appended decision trees (those without pretraining). The framework of RF self-calibration is presented in Fig. 4.

**Figure 4.**
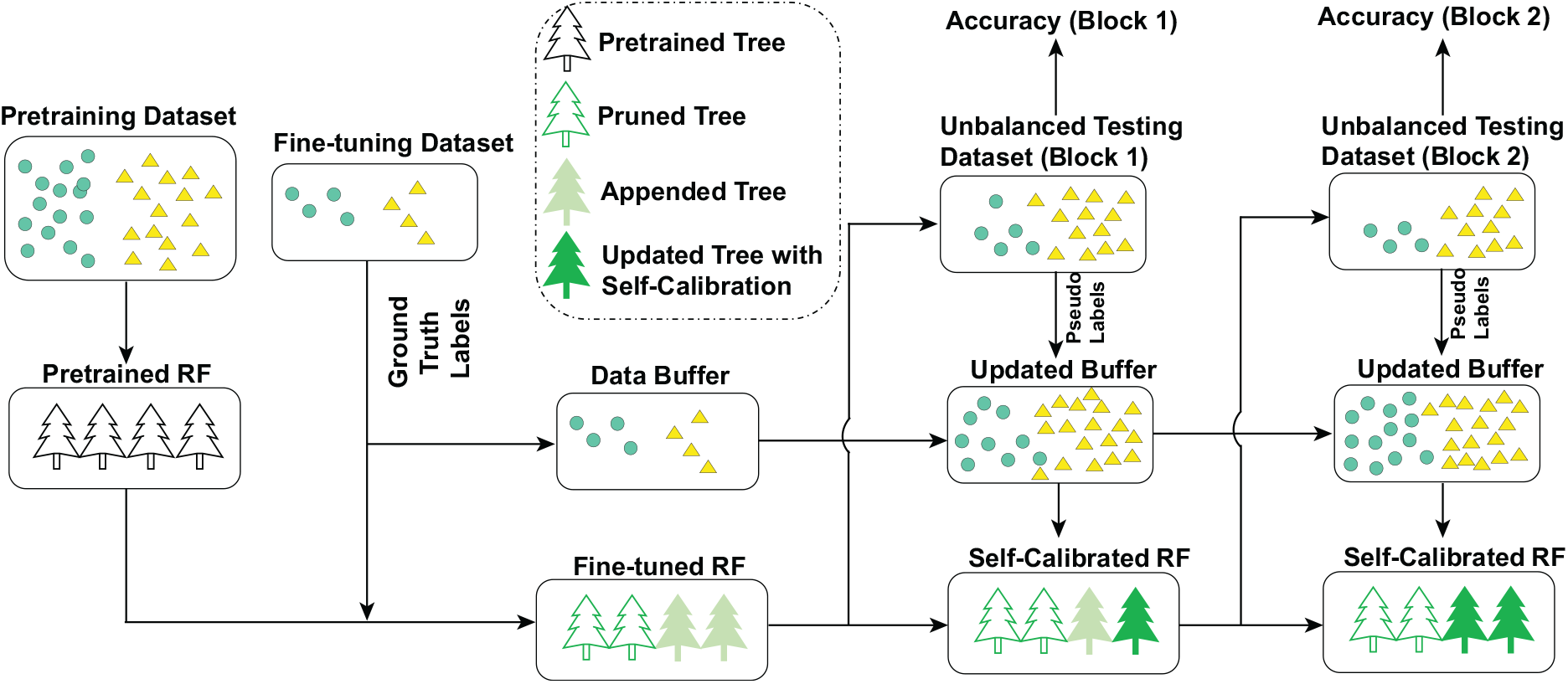
Framework of RF self-calibration.

### 3.5. Evaluation Metrics

The scalability analyses involve evaluating the accuracy of the classification and the total size of the RF model. The classification accuracy was evaluated on each sliding window rather than a complete trial. As for the evaluation of the size of RF models, each decision node of a decision tree consists of four parameters, that is, the index of the left child (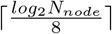 bytes), the index of the right child (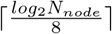 bytes), the index of the selected feature in a node (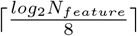 bytes), and the threshold of the selected feature (4 bytes).

### 3.6. Data Allocation

LD datasets 2–4 (66 participants) all include a separate calibration session with 1 trial per hand gesture, enabling the evaluation on the performance of one-shot model fine-tuning. Consequently, when evaluating the LD EMG datasets, each of these 66 participants was viewed as the target test participant one by one, with the other 65 participants and the 20 participants in LD dataset 1 assigned to the pretraining dataset. Data collected in the calibration session of the testing participant were used to fine-tune the pre-trained RF model. All data collected in the testing session of the testing participant were used to evaluate classification accuracy. Self-calibration was performed after each testing block. The performance of RF pre-trained, RF pre-trained and fine-tuned, and self-calibrating RF were compared. Standard linear discriminant analysis (LDA) and standard RF models were implemented to provide baseline performance. Both the standard LDA and the standard RF were trained from scratch on the data collected during the calibration session from the testing participant. Accuracies in multiple test blocks were averaged to evaluate the overall performance of the model.

When evaluating on the HD EMG dataset, a similar leave-one-participant-out strategy was performed, with data from 19 participants in the pretraining set and data from the held out participant in the testing set. The first trial of each hand gesture was assigned to the one-shot calibration set. Considering that the HD data set was not collected in multiple testing blocks (multiple repetitions of each hand gesture were collected in successive trials), the progressive unsupervised self-calibration was not applied in this data set. Only pre-training and supervised fine-tuning of the RF were considered. The pretrained model was fine-tuned only on data in day 1 (the first trial of each hand gesture) and directly applied in day 2 without recalibration. Accuracies on both days were averaged to evaluate the overall performance of the model.

### 3.7. Scalability Analyses

*Scalability with the size of pretraining dataset* The capacity of a model to learn new knowledge from a growing training dataset is essential to examine the upper limit of model performance. Here, we progressively include more participants (from 1 to 85 participants, with an increment of 4) in the pre-training data set (randomly selected) for each testing participant during the leave-one-participant-out cross-validation.

#### 3.7.2 Scalability with the number decision trees

The number of decision trees also has a large impact on the performance and complexity of RF models. We progressively tuned the number of pre-trained decision trees to 200, 150, 100, 75, 50, 80, 30, 20, 10, and 5. In all subsequent sections of the manuscript, the number of decision trees refers to those pre-trained trees. If decision tree appending was applied, the number of appended decision trees was always set the same as the pre-trained trees.

#### 3.7.3. Scalability with the maximal depth of decision trees

Considering that the size of the pretraining dataset is relatively large, the depth of a pre-trained decision tree (the length of decision path from the root node to a leaf node) can be extremely high. By setting the maximal depth of the pre-trained decision trees, we can simplify the decision rules, which in turn affect both the decoding accuracy and the number of parameters. The maximum depth was set to: infinity (no restrictions), 50, 40, 30, 20, 18, 16, 14, 12, 10, 8, 6, and 4.

#### 3.7.4. Scalability with the number of bootstrapping samples

By selecting only a subset of pretraining samples to pre-train each decision tree, the number of nodes in a tree is expected to decrease substantially. Additionally, because different pretrained trees were built on different subsets of pretraining samples, the diversity among decision trees can be improved, contributing to better performance when integrating all trees in the final RF model. The proportion of bootstrapping samples was set to 100%, 50%, 20%, 10%, 5%, 2%, 1%, 0.5%, 0.2%, 0.1%, and 0.05%.

#### 3.7.5. Scalability with the number of electrodes

All of the above analyzes were performed on an LD dataset with 86 participants. We further examine the scalability of an RF model with different numbers of electrodes using the HD dataset. With a total of 256 electrodes in the HD dataset, we evaluated the model characteristics using 256, 224, 192, 160, 128, 96, 64, 32, 16, 8, 4, and 2 electrodes.

### 3.8. Statistical Analyses

For the comparison between ≥ 3 groups, the Friedman test was performed to verify overall significance, followed by the Nemenyi post hoc test for pairwise comparisons. Significance was claimed if *p <* 0.05 was observed.

## 4. Results

### 4.1. Overall Performance Comparison between Different Models

Fig. 5 presents the comparison between the performances of different models, evaluated on LD datasets. By applying decision pruning / grafting, appending, and self-calibration, the accuracy of RF improves progressively. The accuracy of RF with self-calibration significantly outperforms all other models without self-calibration. Note that grafting and pruning operations did not achieve significantly different accuracy. Consequently, only decision tree pruning was performed in our subsequent analyzes, because pruning contributes to a few model parameters rather than grafting by definition.

**Figure 5.**
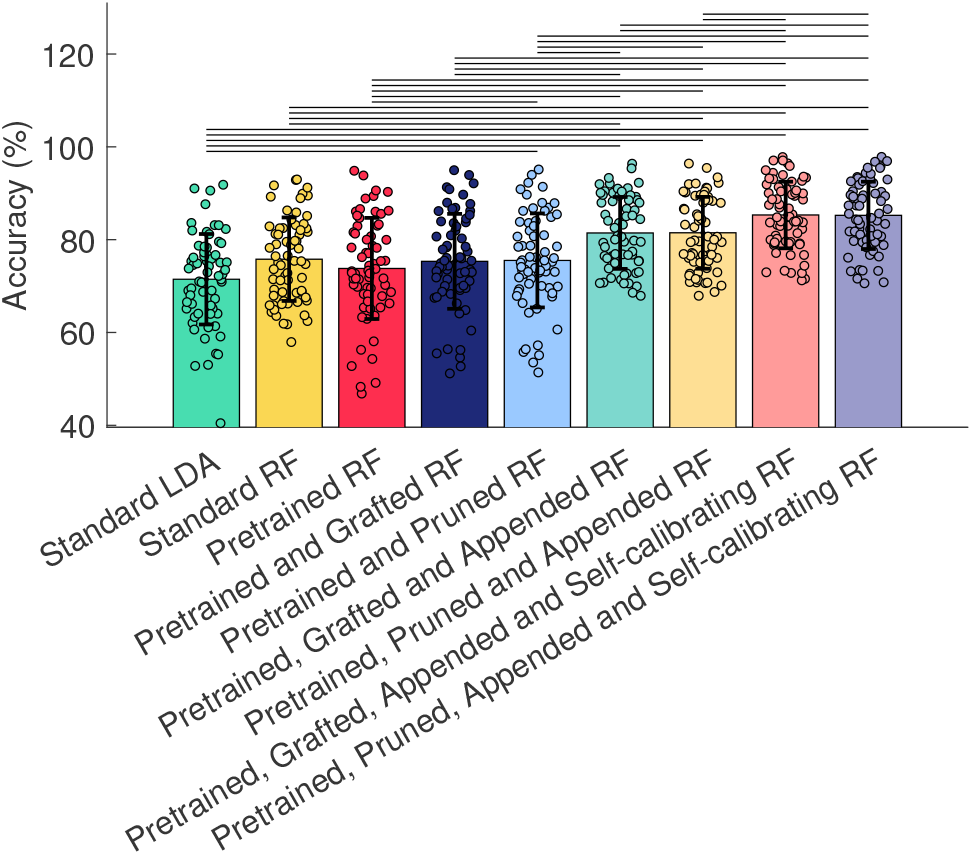
Comparison between different models. The pairs of models with statistically significant difference are marked by horizontal lines. We omitted the * symbols to mark significance for better visualization.

### 4.2. The Effects of Different Factors on Accuracy

Fig. 6 presents the accuracy variation (evaluated in LD datasets) with (1) the number of participants in the pre-training data set (Fig. 6 (a)), (2) the number of decision trees (Fig. 6 (b)), (3) the maximum depth of decision trees (Fig. 6 (c)) and (4) the number of bootstrapping samples to pre-train each decision tree (Fig. 6(d)). Among all variants of RF, the pre-trained RF without any calibration shows the lowest robustness with different conditions of each factor. All variants of RF models show higher robustness on the maximal depth of trees and the number of bootstrapping samples, compared to the number of participants and the number of trees. The number of participants in the pre-training data set contributes to the largest performance variability on pre-trained RF, while the variability substantially decreases when pruning, appending, and self-calibration are applied.

**Figure 6.**
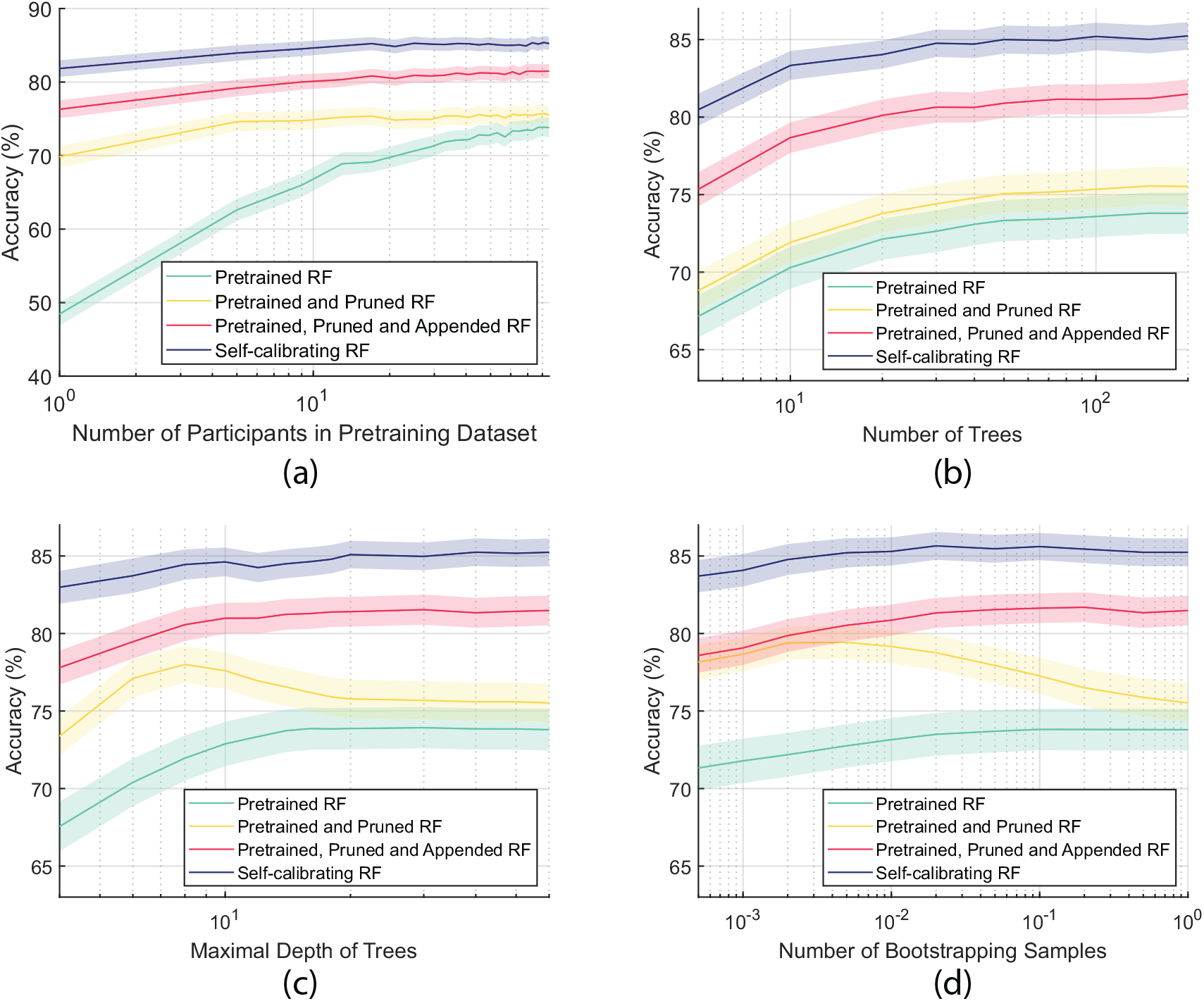
Variation of accuracy with different hyper-parameters. (a) Variation of accuracy with the number of participants in the pretraining dataset. (b) Variation of accuracy with the number of decision trees. (c) Variation of accuracy with the maximal depth of decision trees. (d) Variation of accuracy with the number of bootstrapping samples. For clear visualization, shaded areas indicate the standard error, rather than standard deviation (same for all following figures).

### 4.3. The Variation of Accuracy with Model Size

When tuning each factor presented in Fig. 6, the model size would also vary jointly with the classification accuracy. We therefore present the relation between the accuracy and the model size of different models when tuning different factors in Fig. 7 (evaluated in the LD datasets). According to Fig. 7, the tuning of the number of bootstrapping samples and the tuning of the maximal depth of the decision trees contribute to the most robust model performance with the model size decreasing. Especially in Fig. 7(b), by reducing the number of bootstrapping samples and the maximum tree depth, the classification accuracy characterizes an “arch curve”, which even first increases to the highest accuracy with the model size decreasing. These results demonstrate that simplifying an RF model does not necessarily lead to a reduction in accuracy. By contrast, randomly bootstrapping a small sample set to pre-train each decision tree encourages diversity among all trees. Setting a relatively lower maximum depth for pre-training decision trees can also mitigate overfitting risks. Moreover, pruning a simplified pre-trained decision tree may yield more significant effects. All of these explanations jointly contribute to the “arch curve” in Fig. 7(b). Furthermore, in Fig. 7(d), when tuning the number of bootstrapping samples, the size of the self-calibrating RF decreased from 78.9 MB to 168.0 KB (reduced by 481 ×) with precision reduced from 85. 2% to 83. 7% (reduced by only 1.5%). In all, when a compact RF model was desired in mobile applications, tuning the number of bootstrapping samples or the maximal depth of each tree might be better options.

**Figure 7.**
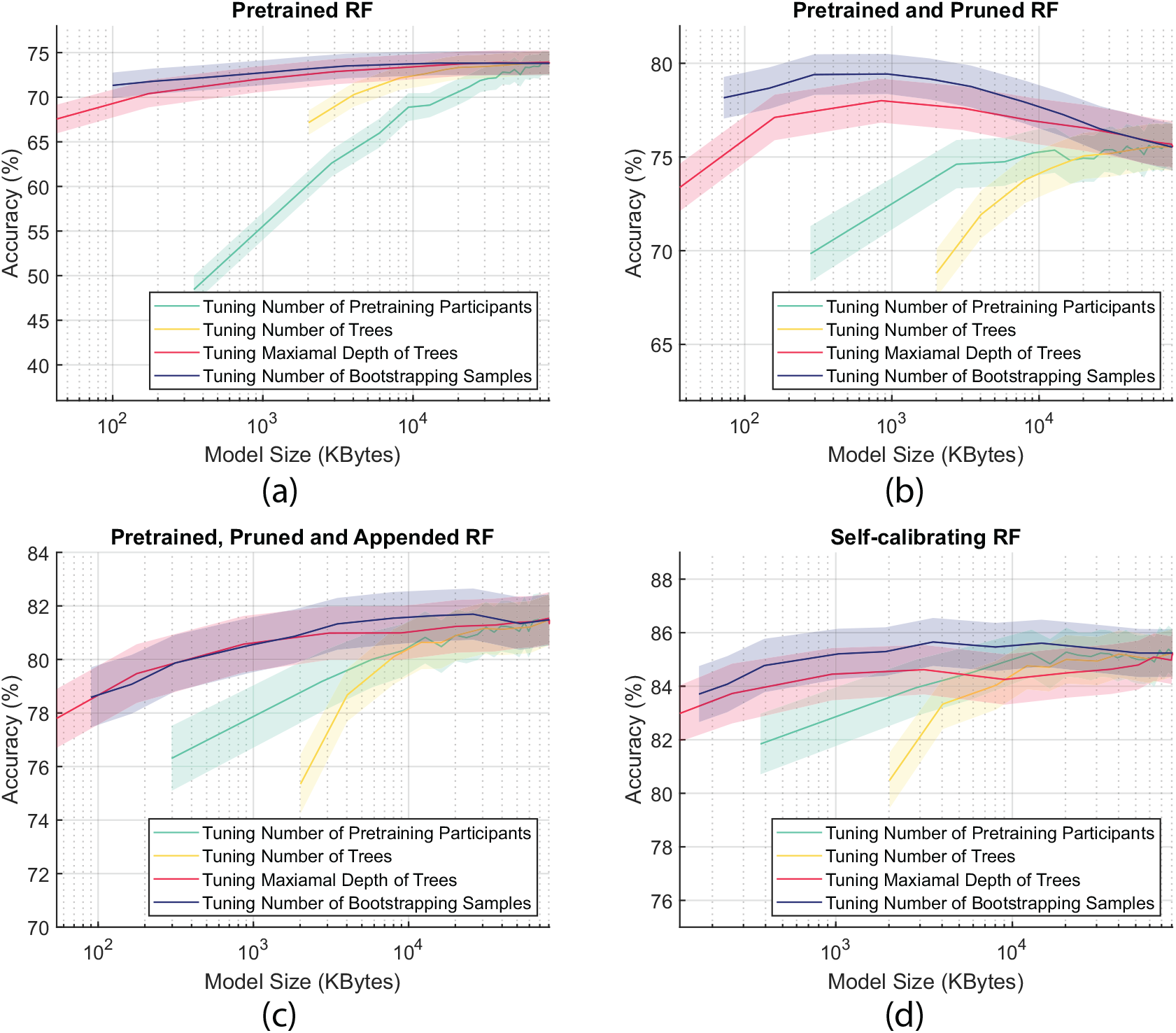
Variation of accuracy with model size. (a) Variation of accuracy (pretrained RF) with model size. (b) Variation of accuracy (pretrained and pruned RF) with model size. (c) Variation of accuracy (pretrained, pruned and appended RF) with model size. (d) Variation of accuracy (self-calibrating RF) with model size.

### 4.4. The Scalability of RF on Electrode Density

Adjusting the number of electrodes in a 256-channel HD EMG affects both the classification accuracy and the size of the model. Fig. 8 shows the relationship between the accuracy of the classification, the number of electrodes, and the size of the model. According to Fig. 8(a), increasing the number of electrodes unsurprisingly improves the accuracy. However, Fig. 8(b) shows that with an increasing number of electrodes, the model size substantially reduces, in contrast to most other machine learning models, which show that a higher-dimensional input normally leads to a model with higher complexity (more parameters). This property of RF can be explained by the fact that, with more alternative features in more available electrodes, each decision tree is more likely to pick more powerful features in each decision node, contributing to a shorter decision path with fewer nodes (parameters). Similar results are illustrated in Fig. 8(c), which shows that a model with a smaller model size (more electrodes) can also achieve higher accuracy. This unique property of RF to a large extent avoids the dilemma of compromising model accuracy for lower model complexity.

**Figure 8.**
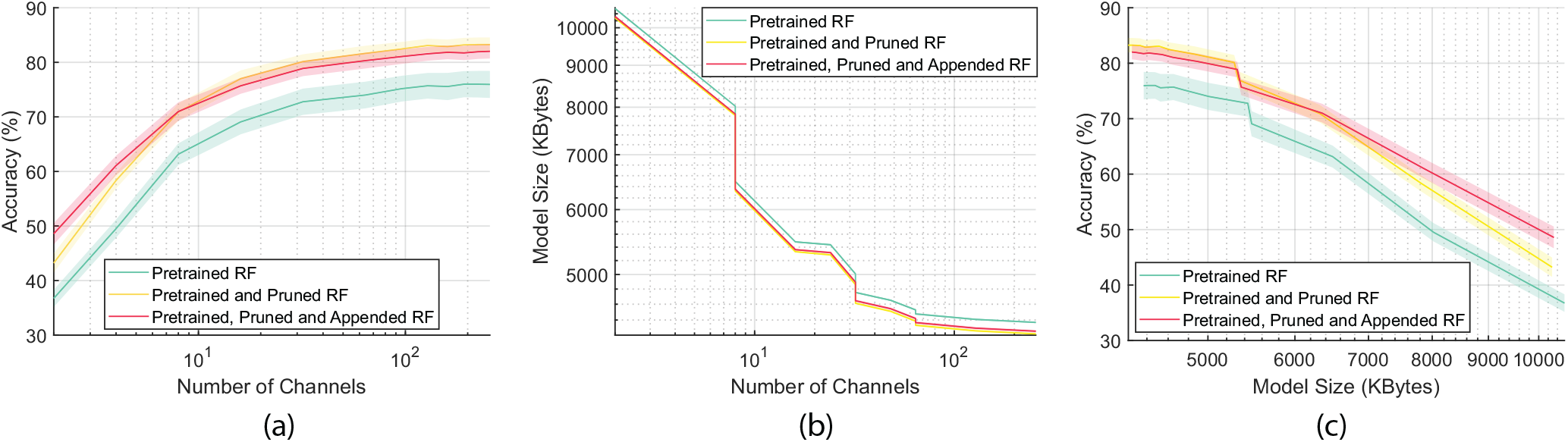
Relation between accuracy, number of electrodes, and model size. (a) Variation of accuracy with the number of electrodes. (b) Variation of model size with the number of electrodes. (c) Variation of accuracy with model size.

Note that in both Fig. 8(b) and Fig. 8(c), the curve shows zigzag patterns. As mentioned above, the number of decision nodes in a decision tree tends to decrease with more electrodes. However, the size of the parameters in each decision node is not constant. For example, when the number of electrodes increases from 16 to 32, the total number of features increases from 160 to 320 (10 features per channel). Therefore, the size of the parameter required to save the index of features in each decision node increases from 1 byte (8 bits, saving the index of up to 2^8^ = 256 features) to 2 bytes (16 bits). Consequently, with the number of electrodes changing from 16 to 32, the number of nodes decreases while the size of parameters in each node increases, leading to the observed zigzag patterns at specific critical values. In general, more electrodes still help reduce the total size of RF models.

## 5. Discussion

Scalability is one of the most important properties of a machine learning model, usually determining the upper limit of the model’s capabilities. Pre-training is a crucial requirement to achieve scalable machine learning. Statistical machine learning models other than neural networks are in general considered capable of being trained only from scratch. By showing RF is pre-trainable, we demonstrate that RF satisfies one important necessary condition to achieve scalable machine learning. We further show that a pre-trained RF model can be one-shot calibrated (for personalisation) on a new user. Such a personalisation process is completely source-free, that is, once the base pre-trained model is built, no access to the source pre-trained model is required, so as to address the privacy issue of biomedical data. Additionally, the RF model can self-calibrate, progressively tracking the dynamic but slowly changing EMG data distribution. Such a pre-trainable, source-free personalisable, adaptable, and at the same time scalable RF model shows high potential in future myoelectric control applications.

Another important condition for scalable machine learning is that the training computational cost is expected to not grow exponentially with the increasing scale of the training data size. In our work, we demonstrate that with only a small proportion of data to train each decision tree, the model can achieve satisfactory and, in certain cases, even better performance. The growing size of training data can contribute to the overall better model performance by improving the diversity of different decision trees trained on different subsets of data, without increasing the training cost of each decision tree. Furthermore, the random forest training is easily parallelisable across different independent decision trees, further improving the practicality in real-world large-scale model training. Scalable machine learning usually faces the common challenge that with a growing size of training dataset, the highly complex model with an extremely large number of samples is difficult to implement on mobile computing devices. We examined different methods to reduce the size of an RF model without compromising the decoding accuracy. Among all these methods, tuning the number of bootstrapping samples and tuning the maximal depth of decision trees are two of the most promising solutions. By tuning the number of bootstrapping samples, the size of the self-calibrating RF could be reduced by ≈ 500 ×, with the accuracy reduced by only 1.5%. All of these findings pave the way for the development of a compact but powerful RF model in mobile myoelectric control applications.

Moreover, the most attractive merit of the RF model reported in this work is the reduction of the model size with a higher dimension of input (more electrodes). Generally, more modalities or more channels of the same modality can provide a substantially larger amount of information, which is a fundamental way to improve model performance. However, a higher dimension of input usually largely increases the number of parameters, imposing challenges on constrained computing devices. For the implementation of most machine learning models, the trade-off between input dimension and model complexity is an essential factor to determine when designing a processing pipeline. The “more is less” property (more electrodes, less parameters) to a large extent avoids the above dilemma. In terms of computational time, the addition of more electrodes would increase the time required for centralized processing of EMG recordings (filtering and feature extraction) in all channels. However, modern digital EMG sensors [28] enable distributed signal processing on electrodes, so the computational time for processing a large number of channels would be constant with a varying number of electrodes. In addition, the use of HD EMG sensors leads to another challenge of channel corruption. With a large number of channels, it is quite common for signals in certain corrupted channels to be contaminated by noise. Our previous study also proved the excellent robustness of RF-based models against noise.

In summary, the RF model shows below merits: (1) is robust to small sample sizes [29], (2) can capture nonlinear relations between samples, (3) is explainable [23], (4) is easily parallelisable and computational efficient, (5) is robust to outliers and noises [23], (6) is pre-trainable, (7) can be one-shot calibrated, (8) can be progressively updated via unsupervised self-calibration, (9) is robust in long-term use [25], (10) is generalisable across body postures/arm positions [13] (11) can be scaled up with a larger dataset, and (12) can be scaled down to reduce model size without largely compromising accuracy. All these merits together unlock the full potential of RF in real-world myoelectric control applications.

## 6. Conclusions

In this work, we systematically explored the scalability of the RF model in myoelectric control. We demonstrate multiple factors to scale up and down an RF model during the process of pretraining, fine-tuning, and automatic self-calibration. Our study explores the effects of various factors such as bootstrapping, decision tree pretraining/pruning/grafting/appending, and the size of the training data. We analyze multiple solutions to trade off between model size and accuracy, with the optimal solution compressing the size of RF models by approximately 500× while decreasing the accuracy by only 1.5%. These findings demonstrate the promising prospects of RF as an alternative foundation model in myoelectric control applications.

## 6.1. Acknowledgments

This work was supported by a grant from Engineering and Physical Sciences Research Council (EPSRC), UK (Grant No. EP/R004242/2).

